# HGF-induced activation of NEPHRIN and NEPH1 serves as a novel mechanism for recovery of podocytes from injury

**DOI:** 10.1101/2020.05.04.077941

**Authors:** Ashish K. Solanki, Pankaj Srivastava, Ehtesham Arif, Christopher M. Furcht, Bushra Rahman, Pei Wen, Avinash Singh, Lawrence B Holzman, Wayne R. Fitzgibbon, Glenn Lobo, Joshua H. Lipschutz, Sang-Ho Kwon, Zhe Han, Matthew J Lazzara, Deepak Nihalani

**Affiliations:** Department of Medicine, Medical University of South Carolina, Charleston, SC, USA; Department of Chemical Engineering, School of Engineering and Applied Science, University of Virginia, Virginia, USA; Department of Medicine, University of Maryland School of Medicine, Baltimore, MD, USA; Biochemistry and Molecular Biology, Medical University of South Carolina, Charleston, SC, USA; Renal-Electrolyte and Hypertension Division, Department of Medicine, University of Pennsylvania, Philadelphia, Pennsylvania, USA; Department of Medicine, Ralph H. Johnson Veterans Affairs Medical Center, Charleston SC, USA; Department of Cellular Biology and Anatomy, Augusta University, Augusta, GA, USA

**Keywords:** HGF, Phosphorylation-dephosphorylation, SHP-2, podocytes

## Abstract

When activated, slit diaphragm proteins NEPHRIN and NEPH1 enable signaling pathways leading to podocyte actin cytoskeleton reorganization, which is critical for podocyte recovery from injury. However, the mechanisms through which these proteins are activated remain unknown. This study presents a novel concept showing ligand-induced activation of NEPHRIN and NEPH1. We first identified phosphatase SHP-2, which directly dephosphorylated these proteins. We next identified HGF, a known SHP-2 modulator, as a rapid inducer of NEPHRIN and NEPH1 phosphorylation. Using baculovirus expressed recombinant purified proteins, SPR (surface plasma resonance), molecular modeling and peptide binding approaches, we show that HGF directly binds NEPHRIN and NEPH1 extracellular domains. Further, using cultured podocytes and Drosophila nephrocytes, we demonstrate that while HGF treatment repaired injured podocytes, the addition of inhibitory NEPH1 or NEPHRIN peptides blocked HGF-induced recovery. Overall, this study shows novel activation and deactivation mechanisms for NEPHRIN and NEPH1 that are required for their function.

## INTRODUCTION

With our increasing knowledge of podocyte biology, it is becoming clear that the glomerular filtration function relies heavily on a properly functioning podocyte. While many proteins contribute to maintain podocyte function, NEPHRIN and NEPH1 are the key proteins that constitute the building blocks of slit diaphragm and are critical for podocyte stability and integrity [1]. Importantly, mutations or genetic deletion of these proteins induce lethality due to severe loss of renal filtration function [2-4]. While many studies demonstrate that NEPHRIN and NEPH1 extracellular domains primarily have a structural function, their intracellular domains are known to assemble a signaling cascade that initiates downstream signaling events leading to actin cytoskeletal changes in podocytes [5]. This implies that these proteins undergo activation events that propagate into downstream signaling. However, the mechanisms behind activation of these proteins remain unknown. Moreover, without the knowledge of a specific ligand that may activate these proteins, the primary function assigned to their extracellular domains remains structural organization of slit diaphragm. In this study, we identified an activation mechanism, where the engagement of their extracellular domains with a ligand HGF induces their phosphorylation. Additionally, the study presents deactivation mechanism, which involves SHP-2-mediated dephosphorylation of these proteins. Collectively, this study provides a greater understanding of how these proteins participate in podocyte development and function.

The cell adhesion proteins NEPHRIN and NEPH1 are known to transduce signals in a Src family kinase Fyn-mediated tyrosine phosphorylation-dependent manner. In podocytes, NEPH1 and NEPHRIN phosphorylation is a proximal event that occurs both, during development and following podocyte injury [6]. The NEPH1 and NEPHRIN phosphorylation-dependent signaling events are primarily involved in regulation of actin dynamics and lamellipodium formation [7, 8] and thereby are critical for structural integrity of slit diaphragm.

One of the major obstacles in investigating the significance of NEPH1 and NEPHRIN phosphorylation following injury has been our lack of understanding of the molecular mechanisms that regulate their phosphorylation. Additionally, the deactivating mechanisms that may involve phosphatases and are critical for maintaining phosphorylation homeostasis, remain unknown.

SHP-2 is a phosphatase that facilitates positive signaling to multiple cellular pathways that control proliferation, migration, differentiation, growth, and survival. SHP-2 binds receptors and adaptor proteins in the presence of extracellular stimuli, including certain cytokines and growth factors including, epidermal growth factor (EGF) [9] and hepatocyte growth factor (HGF) [10]. SHP-2 is involved in regulating the activity of several proteins including Src family kinases, FAK, RhoA/ROCK, and IRS-1 [11]. Although SHP-2 is known to associate with NEPHRIN in a phosphorylation-dependent manner [6], its role as a phosphatase in podocytes remains unclear. While podocyte-specific SHP-2 null mice were resistant to injury [6], it was not clear how its phosphatase function participated in this process. We now present evidences that SHP-2 directly dephosphorylates slit diaphragm proteins NEPHRIN and NEPH1 that is critical for maintaining activation and deactivation homeostasis of these proteins.

Similar to SHP-2, its activator HGF, which is highly elevated in response to injury conditions [12, 13], is also known to induce protective effects in multiple organs, including kidney [14, 15]; however, the mechanism for this protection in podocytes remains unknown. Since our results demonstrate that HGF can induce phosphorylation of NEPHRIN and NEPH1 via a direct interaction, we hypothesize that HGF is the primary regulator of NEPH1 and NEPHRIN signaling, which participates in inducing podocyte recovery from glomerular injury. Accordingly, our novel findings suggest that in response to injury, a recovery response is initiated by HGF in podocytes that involves ligand-based activation of NEPHRIN and NEPH1, leading to actin cytoskeletal reorganization. Overall, we present compelling evidence for the ligand-based activation and deactivation mechanisms for NEPHRIN and NEPH1.

## Results

### SHP-2 is a novel binding partner for NEPH1

To identify NEPH1 binding proteins, we performed mass spectrometry analysis of NEPH1-immunoprecipitated complex from podocytes. SHP-2, a product of the PTPN11 gene, was identified as a novel binding partner. (**Fig. 1A**). Since SHP-2 bound NEPHRIN in a phosphorylation-dependent manner [16], we investigated whether NEPH1 phosphorylation also enhanced SHP-2 binding. Similar to NEPHRIN, the phosphorylation of NEPH1 either via treatment of podocytes with pervanadate **[17-19]**, or by co-expressing with FYN kinase, significantly increased NEPH1 and SHP-2 interaction (**Fig. 1B and C**). By mixing recombinant purified proteins, we further demonstrated a direct interaction between the cytoplasmic domain of NEPH1 and SHP-2 (**Fig. 1D**). To further determine if NEPH1 is a substrate for SHP-2, we tested the binding of NEPH1 with a substrate trapping SHP-2^DM^ mutant **[20, 21]**. This mutant displayed much higher ability to trap phosphorylated NEPH1 than the wild-type SHP-2 (**Suppl. Fig 1**), suggesting a functional interaction between the two proteins.

**Figure 1:**
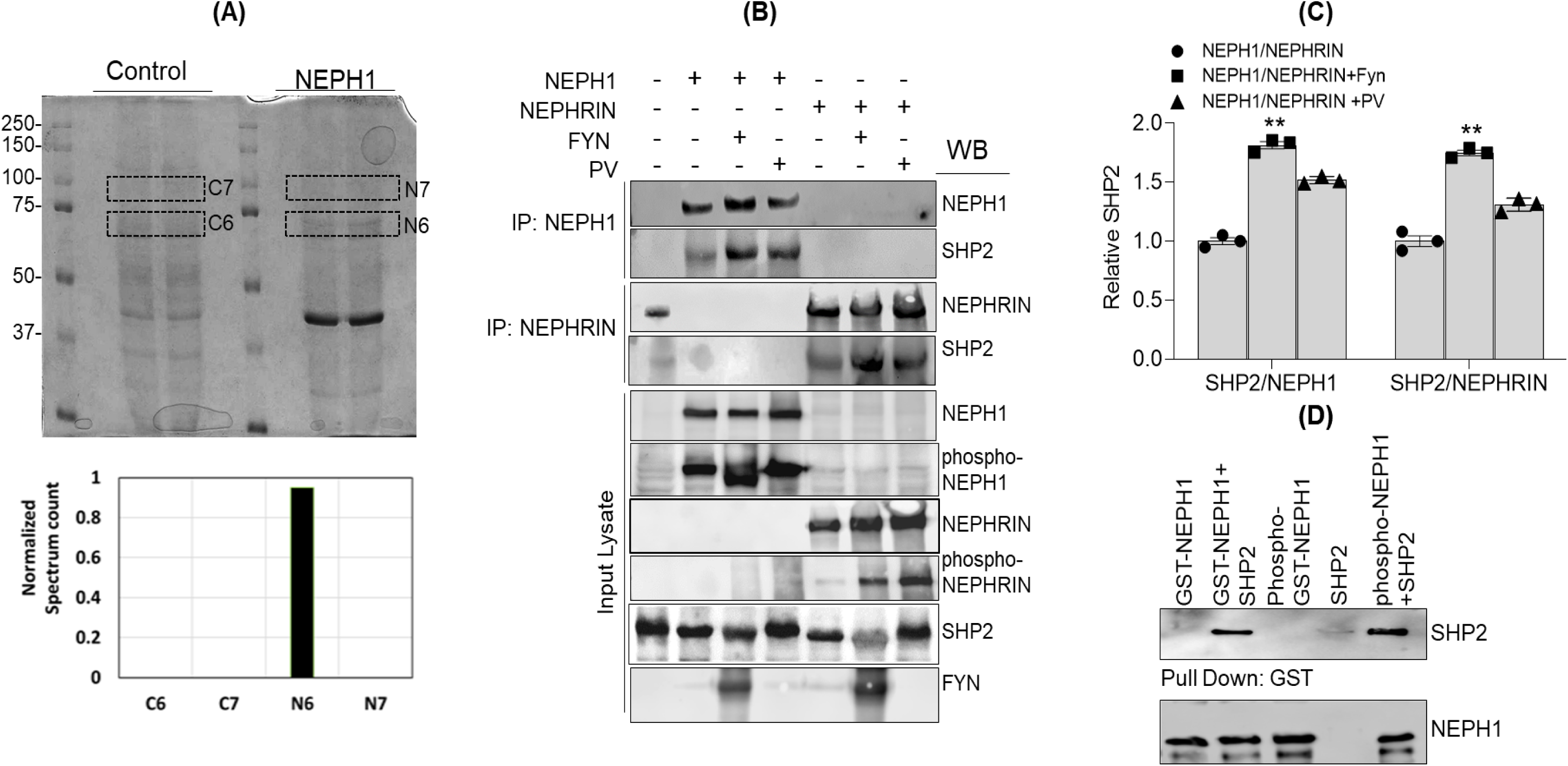
SHP-2 binds NEPH1: **(A)** Immunoprecipitation of the endogenous NEPH1 from podocytes followed by mass spectrometric analysis identified SHP-2 (N6) as a novel NEPH1 interacting protein **(B & C)** NEPH1 or NEPHRIN were phosphorylated by either coexpressing with FYN or PV treatment and were immunoprecipitated with their respective antibodies. Western blotting of the immunoprecipitated complexes showed increased SHP-2 binding to the phosphorylated NEPHRIN and NEPH1. All experiments were performed in triplicate. Data are presented as mean ± SEM, and P values were calculated using the Kruskal–Wallis one-way analysis of variance. **P ≤ 0.005 **(D)** Direct binding was evaluated by mixing purified phosphorylated GST-NEPH1 (produced in TKB1 cells) with purified recombinant SHP-2, which showed increased binding of SHP-2 with phosphorylated GST-NEPH1.

### SHP-2 dephosphorylates NEPHRIN and NEPH1

Although a previous report suggested that co-expression of SHP-2 with NEPHRIN enhanced its phosphorylation due to its effect on FYN activation (28), a direct effect of SHP-2 on NEPHRIN was not evaluated. Since SHP-2 is a phosphatase (31, 32), we hypothesized that SHP-2 directly dephosphorylates NEPH1 and NEPHRIN. To test this, we either co-expressed NEPH1 or NEPHRIN with FYN in HEK293 cells or treated the NEPH1 and NEPHRIN expressing stable cultured podocytes [22] with pervanadate and immunoprecipitated with their respective antibodies. The immunoprecipitated complexes containing the phosphorylated NEPHRIN or NEPH1 were incubated with purified active recombinant SHP-2 in a phosphatase buffer. Interestingly, significant dephosphorylation of both, NEPHRIN and NEPH1 was noted in the presence of SHP-2 (**Fig. 2A-D**). In a subsequent experiment, we immobilized purified cytoplasmic domains of GST-NEPH1 or GST-NEPHRIN on beads and incubated them with purified active recombinant FYN. This resulted in phosphorylation of the two proteins. Following washing, purified active recombinant SHP-2 was added, which indicated dephosphorylation of these proteins in a time-dependent fashion (**Fig. 2E-F**). Collectively, these results demonstrate that SHP-2 is a phosphatase for NEPHRIN and NEPH1.

**Figure 2:**
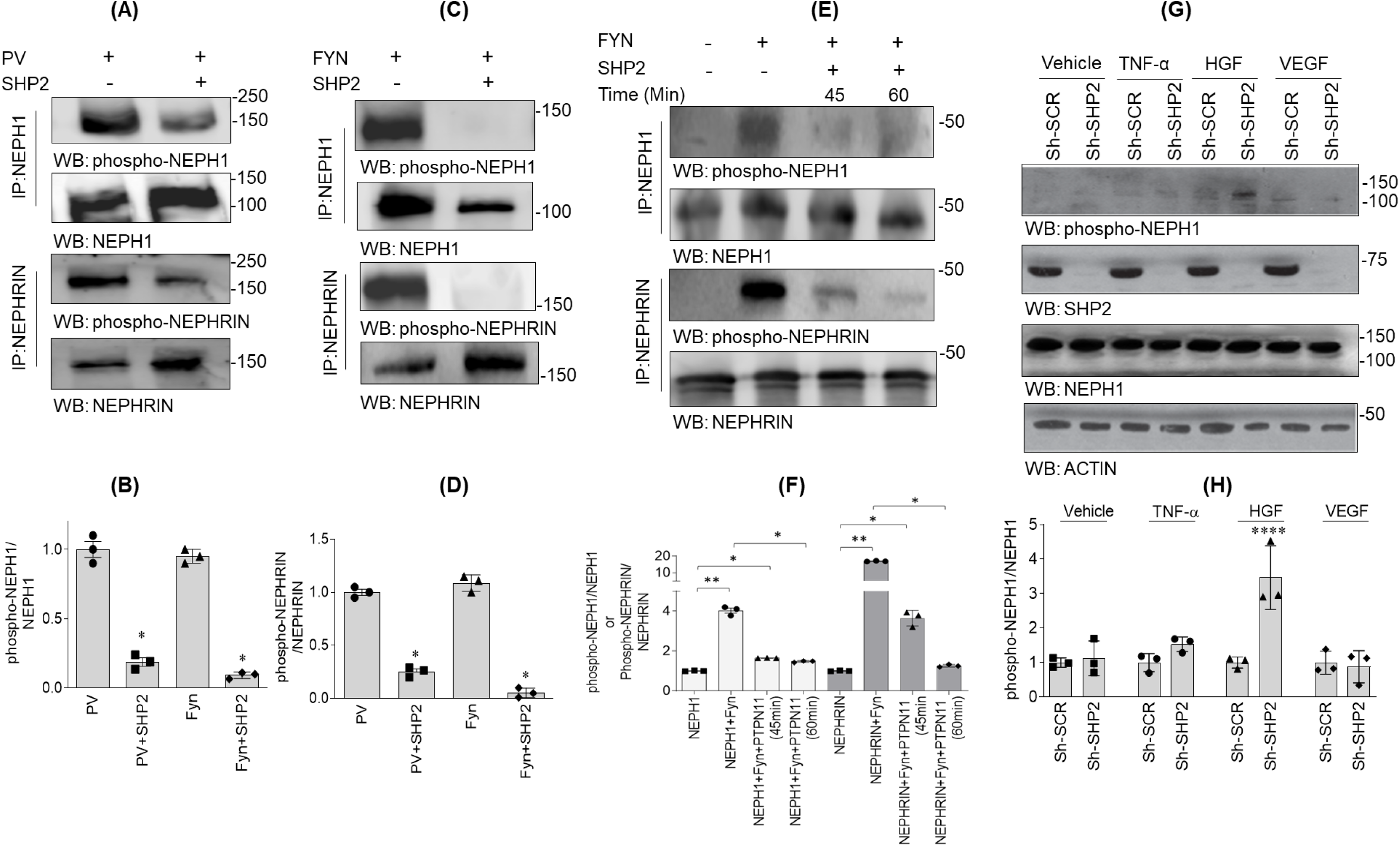
SHP-2 is a phosphatase for NEPH1 and NEPHRIN: **(A & B)** HEK293 cells expressing NEPHRIN or NEPH1 were treated with PV followed by immunoprecipitation of NEPH1 and NEPHRIN. Recombinant SHP-2 protein was added to the immunoprecipitated complexes and the extent of dephosphorylation was measured by respective phospho antibodies. All experiments were performed in triplicate. Data are presented as mean ± SEM, and P values were calculated using the Mann-Whitney (nonparametric) test, one tailed. *P ≤ 0.05 **(C & D)** NEPH1 and NEPHRIN were overexpressed in HEK293 cells and the proteins were immunoprecipitated using their respective antibodies and incubated with purified FYN (500 ng). Following washing with PBST, the phosphorylated proteins were incubated with purified recombinant active SHP-2 and the extent of dephosphorylation was measured by respective phospho antibodies. All experiments were performed in triplicate. Data are presented as mean ± SEM, and P values were calculated using the Mann-Whitney (nonparametric) test, one tailed. *P ≤ 0.05 **(E & F)** Purified recombinant GST-NEPH1CD and GST-NEPHRINCD (cytoplasmic domains) proteins were phosphorylated by incubating with recombinant active FYN immobilized on nickel beads. Post phosphorylation, FYN beads were removed and the phosphorylated GST-NEPH1CD and GST-NEPHRINCD proteins were incubated with purified recombinant SHP-2 for indicated times and western blotted with respective phospho antibodies, which confirmed SHP-2 mediated dephosphorylation. All experiments were performed in triplicate. Data are presented as mean ± SEM, and P values were calculated using the Kruskal–Wallis one-way analysis of variance. *P ≤ 0.05, **P ≤ 0.005. **(G & H)** Control podocytes and podocytes with stable SHP-2 knockdown were treated with various growth factors (including TNF-α, HGF and VEGF) and the phosphorylation of endogenous NEPH1 was evaluated by phospho-NEPH1 antibody. NEPH1 phosphorylation was only visible under HGF stimulation of SHP-2 KD podocytes. SCR=scrambled. All experiments were performed in triplicate. Data are presented as mean ± SEM, and P values were calculated using the Sidak’s multiple comparisons test (two-way anova). ****P ≤0.0001

### HGF is a novel inducer of NEPHRIN and NEPH1 phosphorylation

Since SHP-2 appeared as a potent phosphatase for NEPHRIN and NEPH1, we hypothesized that the ligand-induced phosphorylation of NEPHRIN and NEPH1 may be suppressed in the presence of SHP-2. Thus, to evaluate phosphorylation of these proteins, we constructed stable SHP-2 knockdown (KD) cultured human podocytes that are known to endogenously express NEPH1 [23-26]. These podocytes were screened against various growth factors and the phosphorylation of NEPH1 was evaluated using a phospho NEPH1 antibody [19, 26, 27]. Interestingly, phosphorylation of endogenous NEPH1 was only visible in SHP-2 knockdown podocytes treated with HGF (20ng/ml) (**Fig. 2G-H**), which is (also) a potent activator of MET receptor and SHP-2 [21, 28, 29]. Since this result has huge implications, we performed several additional experiments to confirm if similar effects were observed for NEPHRIN. By co-expressing NEPHRIN and NEPH1 in HEK293 cells with HGF and blotting the cell lysates with respective phospho-antibodies, we first showed that these proteins are phosphorylated in the presence of HGF (**Fig. 3A**). Additionally, we exogenously added purified active recombinant HGF to NEPH1 or NEPHRIN-overexpressing HEK293 cells, which showed a time-dependent increase in their phosphorylation by HGF (**Fig. 3B-C**). Lastly, we performed transwell assay (**Fig. 3D**), where HEK293 cells overexpressing HGF in culture medium were cultured in the upper chamber of transwell plate and NEPHRIN or NEPH1 overexpressing HEK293 cells were cultured in the bottom chamber. Interestingly, significant NEPHRIN and NEPH1 phosphorylations were noted only when the upper trans chamber contained HGF-expressing cells (**Fig. 3E**). Collectively, these results confirmed the role of HGF as a potential ligand and inducer of NEPHRIN and NEPH1 phosphorylation.

**Figure 3:**
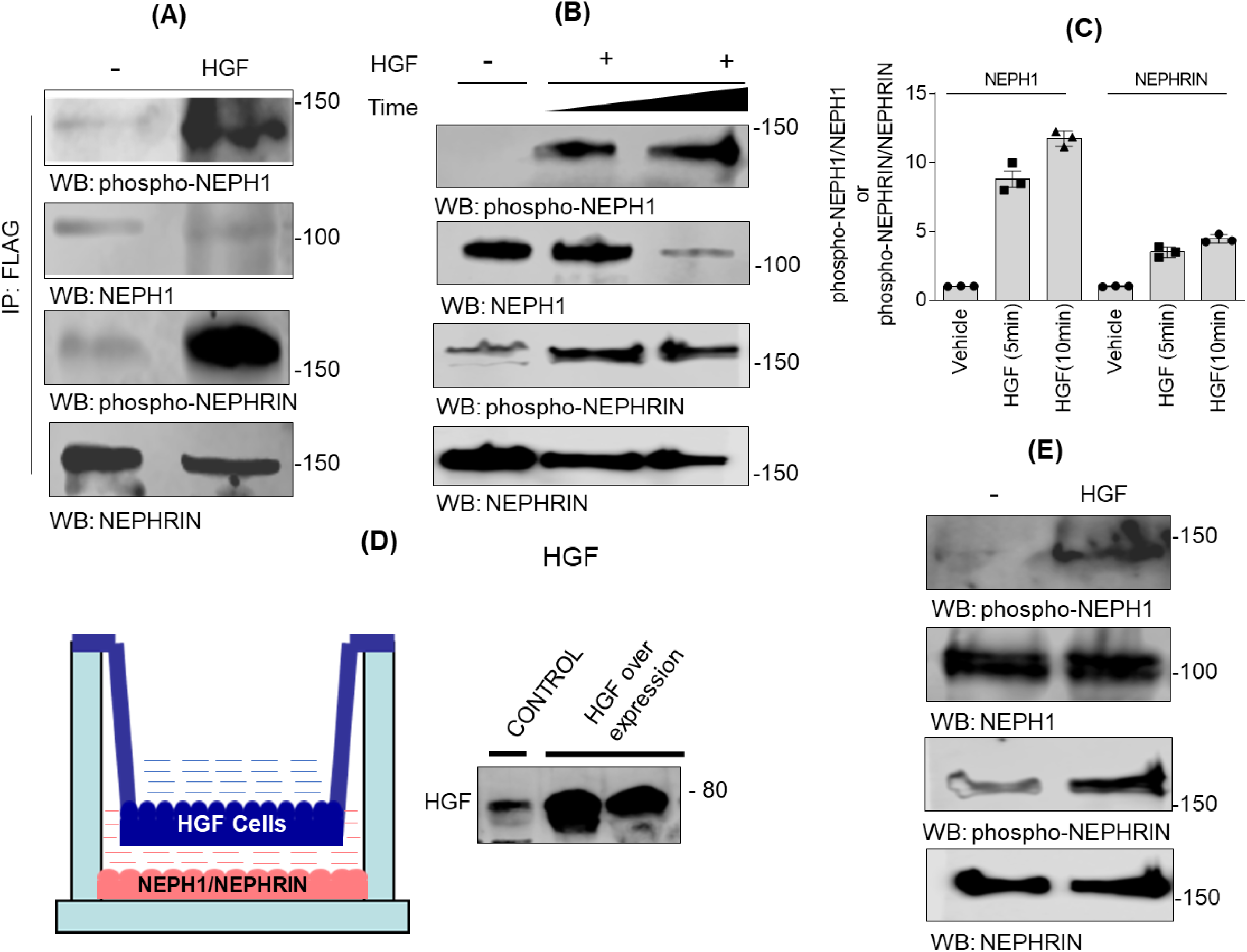
HGF is the novel inducer of NEPHRIN and NEPH1 phosphorylation: **(A)** NEPHRIN and NEPH1 were coexpressed with HGF in HEK293 cells and phosphorylation was analyzed using their respective phospho-antibodies. **(B & C)** Time dependent increase in the phosphorylation of NEPH1 and NEPHRIN was observed when purified recombinant active HGF was exogenously added to the NEPH1 and NEPHRIN overexpressing HEK293 cells. **(D & E)** Schematic of the trans-well plate experimental set up. HGF expressing HEK293 were grown in the upper trans-well chamber and NEPH1 and NEPHRIN expressing stable HEK293 cells were cultured in the lower chamber. NEPHRIN and NEPH1 phosphorylation was measured by lysing the cells in lower chamber and western blotting with their respective phospho antibodies. All experiments were performed in triplicate. Data are presented as mean ± SEM, and P values were calculated using the Kruskal–Wallis one-way analysis of variance. **P ≤ 0.0001

### MET receptor is not required for HGF-induced phosphorylation of NEPHRIN and NEPH1

Since HGF is also a potent activator of MET receptor [12, 13], it is possible that MET may be directly or indirectly involved in NEPHRIN and NEPH1 phosphorylation. To test this, we constructed a stable knockdown of MET receptor in HEK293 cells overexpressing NEPHRIN or NEPH1 and performed similar HGF-induced phosphorylation. The results showed no change in HGF-mediated phosphorylation of NEPHRIN or NEPH1 in MET-knockdown cells (**Suppl. Fig. 2A-B**), suggesting that HGF induces NEPHRIN and NEPH1 phosphorylation independently of the MET receptor. Additionally, we used a potent MET receptor inhibitor (Crizotinib) [30] to further demonstrate that MET is not required for HGF-induced phosphorylation of NEPHRIN and NEPH1 (**Suppl. Fig. 2C**).

### HGF is a novel ligand that binds NEPHRIN and NEPH1 extracellular domains

If HGF acts as a ligand then it should interact extracellularly with NEPHRIN and NEPH1. To test this, we first co-expressed HGF with NEPHRIN or NEPH1 and determined the binding through immunoprecipitation. This showed that both NEPHRIN and NEPH1 interact with HGF (**Fig. 4A**). To determine whether the interaction is direct, we performed two different experiments. First, we used the HGF-binding site of the MET receptor to identify potential binding regions in the extracellular domains of NEPHRIN and NEPH1 through sequence homology and molecular modeling, which indicated IgG3 in NEPH1 and IgG2 in NEPHRIN as HGF interacting domains (**Suppl. Fig. 3)**. Importantly, these regions were highly conserved among various species (**Suppl. Fig. 4**). The peptides from these regions were synthesized and used in a dot blot assay, where peptides were immobilized and probed with active recombinant HGF. The results showed that HGF interacted with NEPH1 peptide and NEPHRIN peptide 1 but not with NEPHRIN peptide 2 (**Fig. 4B**). To further determine a direct interaction, significant effort was spent generating mammalian forms of recombinant NEPHRIN and NEPH1. As shown in **Fig. 4C**, we demonstrate the first report of an SF9 insect cell line expressing mammalian His-FLAG-NEPH1-FL (full length) and His-FLAG-NEPHRIN-ECD (entire extracellular domain), which was established using the Baculoviral Expression system. The purified recombinant NEPHRIN and NEPH1 proteins were mixed with commercially obtained purified and active HGF **(Fig. 4C)** and SPR (surface plasma resonance) was employed to evaluate their binding. The results showed a concentration-dependent direct interaction of HGF with NEPHRIN or NEPH1 **(Fig. 4D-E)**. Interestingly, while the interaction of HGF with NEPH1 was in the nanomolar range (K_D_=25.7nM) (**Table 1a**), the interaction with NEPHRIN was in the picomolar range (K_D_=0.4nM) (**Table 1b**), suggesting that HGF binds NEPHRIN with much higher affinity (**Fig. 4D-E**).

**Figure 4:**
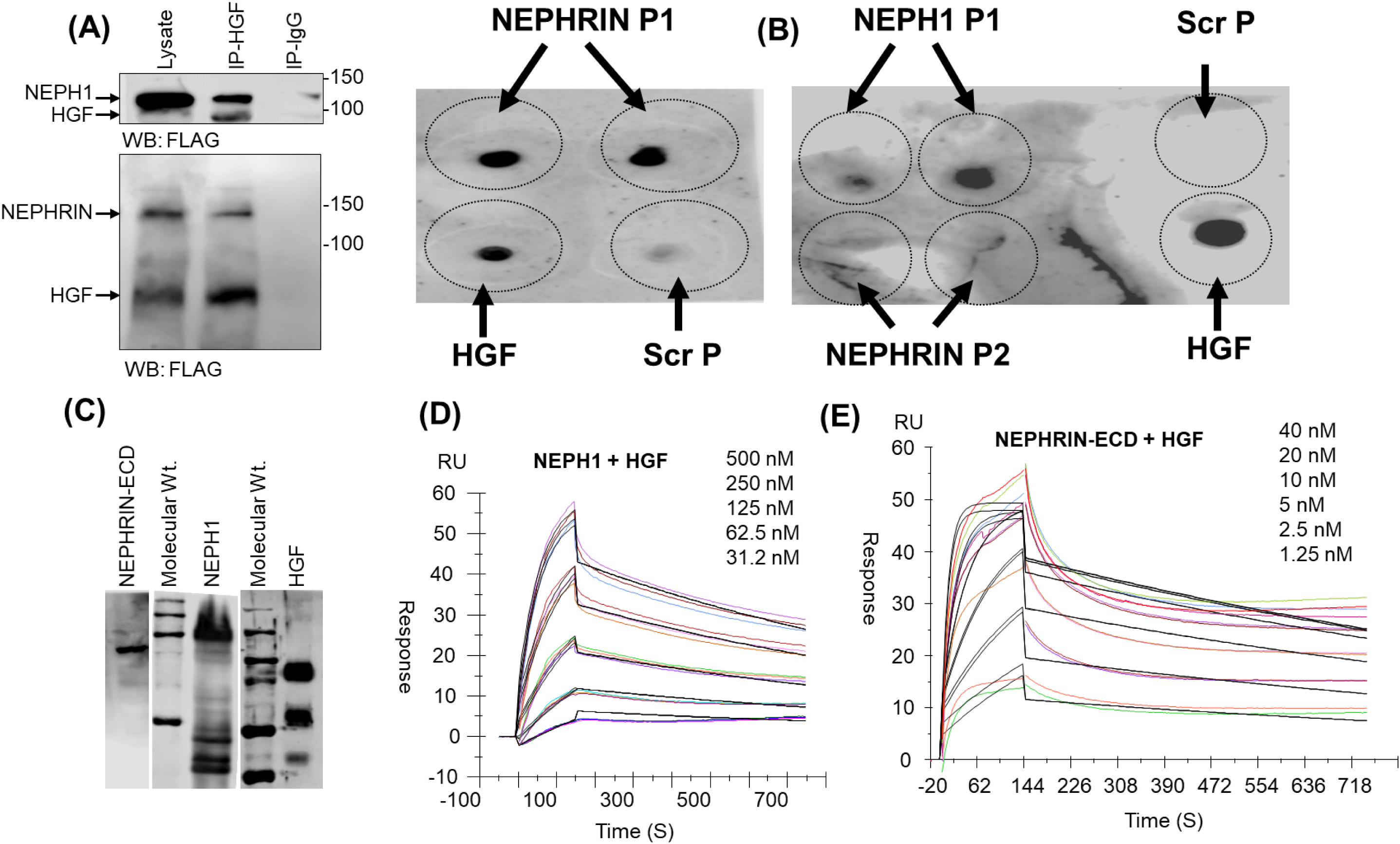
HGF interacts with NEPHRIN and NEPH1: **(A)** NEPHRIN and NEPH1 were coexpressed with HGF in HEK293 cells. HGF was immunoprecipitated using HGF antibody and the immunoprecipitated complexes were analyzed for NEPHRIN and NEPH1 binding by western blotting using FLAG antibody. **(B) Dot Blot assay:** Synthetic peptides corresponding to the HGF binding regions of NEPHRIN and NEPH1 were spotted onto nitrocellulose membrane and dot blot and probed with recombinant HGF. HGF binding was apparent with peptides NEPH1-P1 and NEPHRIN-P1, but not with NEPHRIN-P2 and control scramble (Scr) peptides. **(C-E)** Baculovirus expressed NEPHRIN-ECD (extracellular domain), and NEPH1 (**C**), were mixed with HGF in the indicated amounts and subjected to SPR. SPR analysis showed concentration-dependent binding of NEPH1 (**D**) and NEPHRIN (**E**) with HGF. The calculated KD values for NEPHRIN and NEPH1 are presented in Tables 1a & 1b.

### HGF induces podocytes recovery from injury in the *in vitro* and *in vivo* models of podocyte injury

Since HGF functions as an injury-induced effector for tissue repair, we hypothesized that HGF is involved in podocyte repair following injury. To test this, cultured human podocytes overexpressing NEPH1 were injured with PS (protamine sulphate), as described previously [19, 24, 31, 32], which resulted in severe actin cytoskeleton disorganization **(Fig. 5A-B)**. Interestingly, addition of exogenous recombinant HGF resulted in significant recovery of actin cytoskeleton **(Fig. 5A-B)**. Since the cultured podocytes express measurable amounts of endogenous NEPH1 but not NEPHRIN, addition of the NEPH1 peptide (that binds HGF), or by using NEPH1 knockdown podocytes in this system, blocked HGF-induced recovery (**Fig 5A-B**). In parallel, we performed a similar experiment with NEPHRIN-overexpressing podocytes, which showed similar HGF-induced recovery of podocytes from injury, which was blocked by the NEPHRIN peptide1 (**Suppl. Fig. 6A-B**).

**Figure 5:**
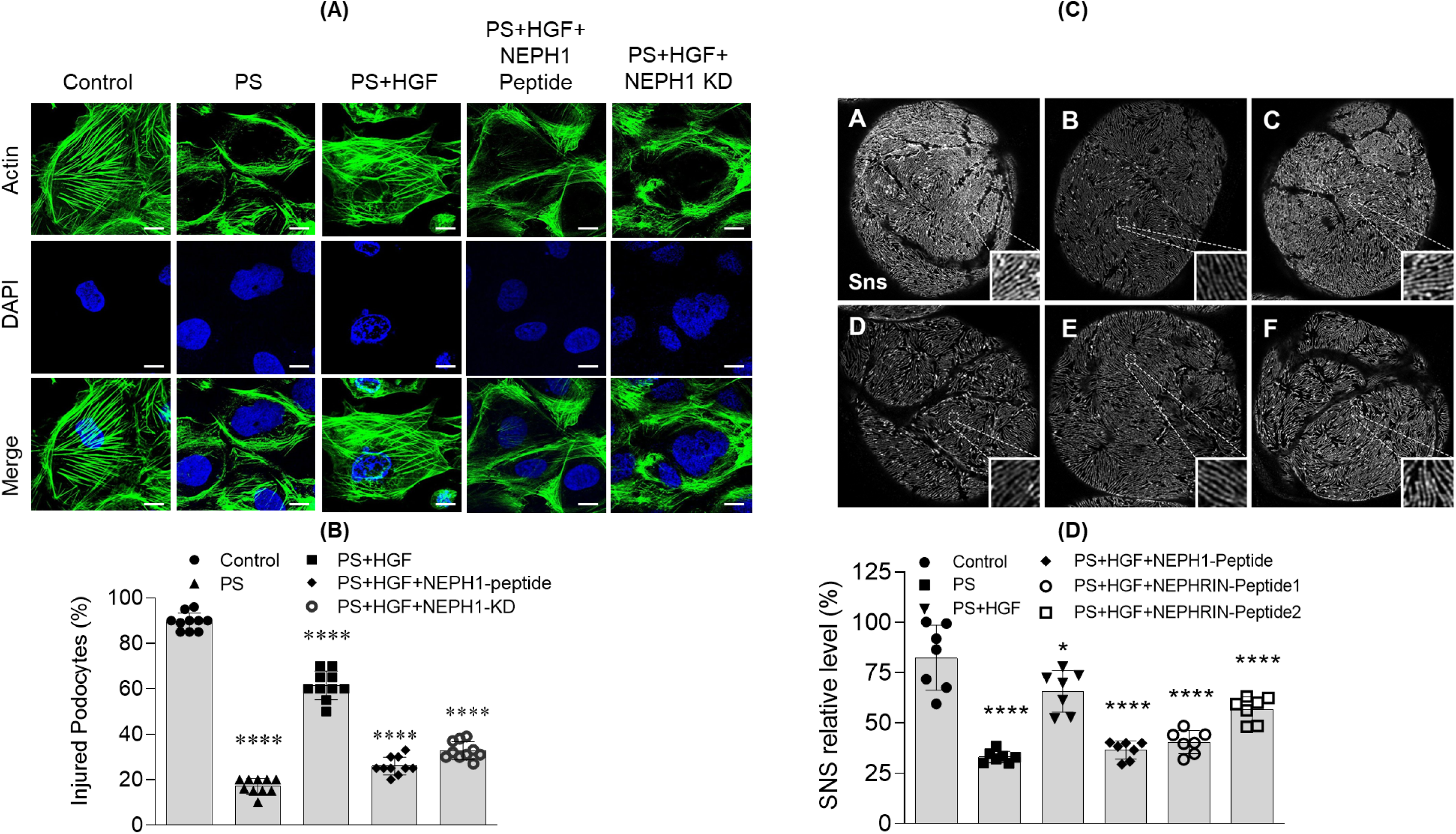
HGF treatment repairs podocytes/nephrocytes in a NEPH1/NEPHRIN dependent fashion: (**A & B**) Cultured human podocytes were treated with Protamine Sulphate (PS) and actin cytoskeleton (green) disorganization was visualized by phalloidin staining. To induce recovery, HGF (50 ng/ml) was added to the PS treated podocytes. Addition of NEPH1 inhibitory peptide blocked HGF-induced recovery. 10 cells per experimental condition were evaluated from three experimental repeats. Scalebar=25µm. Data are presented as mean ± SEM, and P values were calculated using the Tukey’s multiple comparisons test (One-way anova). *P ≤0.01, ****P ≤0.0001 (**C & D**) Sns staining of Drosophila pericardial nephrocytes treated with PS in the absence or presence of HGF and NEPHRIN or NEPH1 peptides. Decreased Sns staining was noted in PS treated nephrocytes, which was rescued by treatment with HGF. Further addition of HGF-interacting NEPH1 P1 and NEPHRIN P1 peptides, but not NEPHRIN P2 peptide, blocked the effect of HGF. For quantification, 7 nephrocytes from three flies for each condition, were analyzed. Data are presented as mean ± SEM, and P values were calculated using the Tukey’s multiple comparisons test (One-way anova). ****P ≤0.0001.

**Figure 6:**
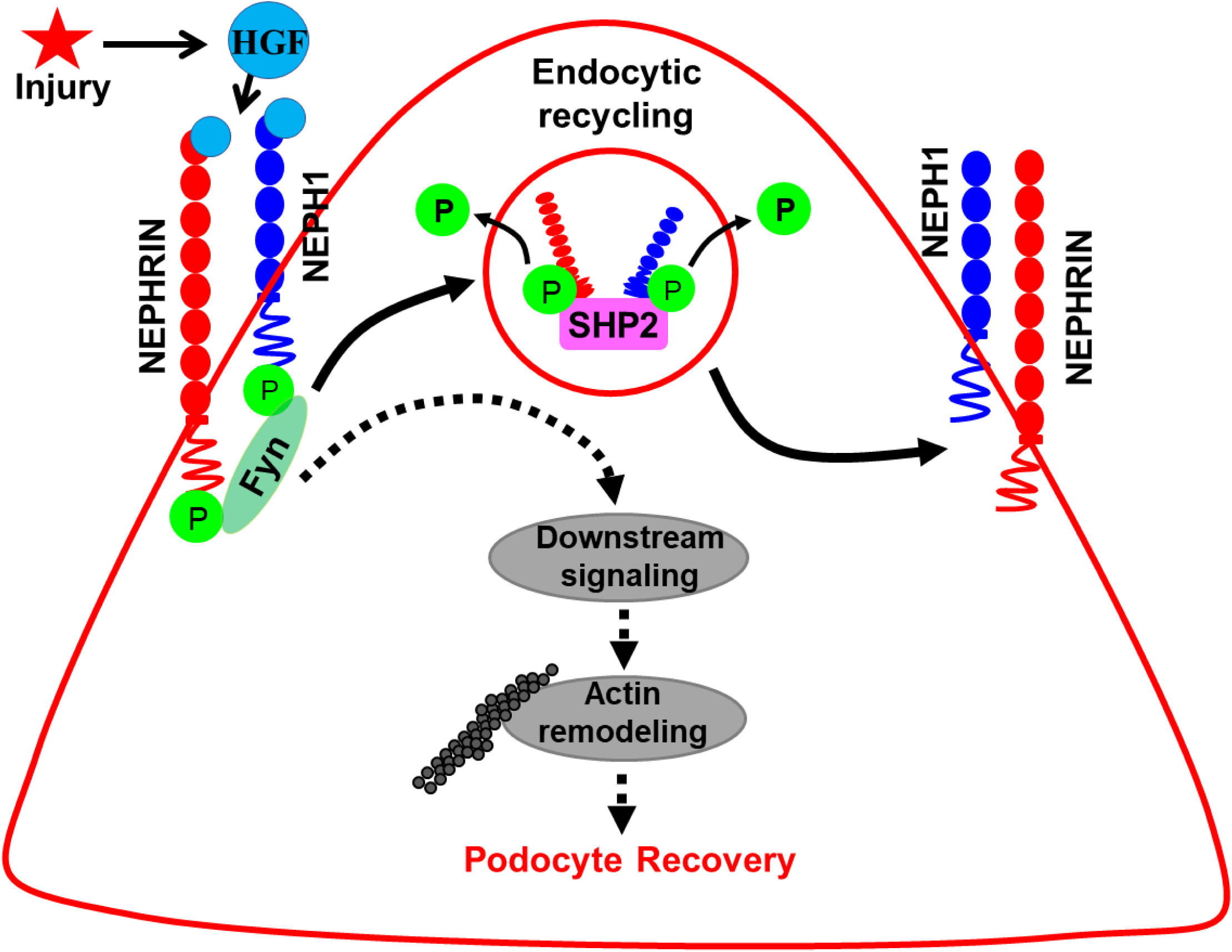
Schematic of the activation and deactivation mechanisms for NEPHRIN and NEPH1 in podocytes. In response to injury, HGF initiates recovery process by activating NEPHRIN and NEPH1, which induces actin cytoskeleton reorganization, leading to podocyte repair. To maintain homeostasis, subsequent deactivation is mediated in a SHP-2 dependent fashion.

To further analyze HGF-induced recovery, Drosophila nephrocytes were used, where nephrocytes isolated from Drosophila were subjected to Protamine-sulphate (PS) treatment and stained with SNS (NEPHRIN) antibody to evaluate the extent of injury [33]. We observed a significant loss of SNS protein, which was rescued by the HGF treatment (**Figure 5C-D**). Furthermore, the HGF-interacting peptides, NEPH1-P1 and NEPHRIN P1, blocked the effect of HGF. In contrast, the NEPHRIN P2, which did not interact with HGF, was unable to block the recovery effect of HGF. These results further emphasize the fact that HGF induces podocyte recovery in a NEPHRIN and NEPH1 dependent fashion.

## Discussion

The filtration function of a glomerulus is highly dependent on podocytes since numerous reports demonstrate that injury to podocytes and loss of their modified tight junction slit diaphragm is directly associated with glomerular damage and loss of renal filtration function [34]. Thus, identification of factors that regulate podocyte repair, are of vital importance in understanding the mechanisms underlying podocyte survival and function and in identifying new therapeutic strategies.

Several proteins are known to participate in maintaining the structural integrity and composition of slit diaphragm including NEPHRIN and NEPH1, whose genetic defects in humans and mice lead to FSGS, which is characterized by podocyte dysfunction and nephrotic range proteinuria [19, 27, 34-38]. The prevailing hypotheses suggest that the extracellular domains of NEPHRIN and NEPH1 constitute the structural framework of slit diaphragm, thus, maintaining its structural integrity [17, 37, 39, 40]. We now present compelling evidences that NEPHRIN and NEPH1 have receptor like properties, where they can be activated in a ligand-based fashion by HGF and are deactivated by the phosphatase SHP-2. Previous biochemical and genetic studies have shown that Src family kinases (SFK’s) mediated tyrosine phosphorylations of NEPHRIN and NEPH1 cytoplasmic domains subsequently induce signal transduction events leading to podocyte actin cytoskeletal reorganization [16, 18, 36, 41]. However, the activating and deactivating mechanisms that may regulate the phosphorylation of these proteins remain unknown [17, 19, 39]. The data presented herein suggest that, SHP-2 acts as a phosphatase that directly dephosphorylates NEPH1 and NEPHRIN. Since injury to podocytes is known to induce NEPHRIN and NEPH1 phosphorylation [6] [7, 8], we hypothesized that such induction may require downregulation of SHP-2 expression. Accordingly, the mRNA profiling of cultured podocytes injured by puromycin amino nucleoside (Gene Expression Omnibus accession number GSE127736, [22] showed five-fold reduction in SHP-2 expression (**Suppl. Fig. 7**). Collectively, these results are consistent with a role for SHP-2 in regulating NEPHRIN and NEPH1 activation. Further, this may highlight a possible protective mechanism observed in SHP-2 KO mice [42], whereby loss of SHP-2 may allow consistent activation of NEPHRIN and NEPH1, leading to efficient repair of podocyte actin cytoskeleton. While this may explain a deactivation mechanism for NEPHRIN and NEPH1, what biochemical stimulus activates these proteins has so far remained a mystery. Strikingly, we observed that HGF that induces SHP-2 activation via MET receptor also induced the phosphorylation of NEPHRIN and NEPH1. We performed several biochemical experiments to establish HGF as the inducer of NEPHRIN and NEPH1 phosphorylation (**Fig. 2G & Fig. 3)**. Since HGF works as an injury-induced effector for tissue repair, acting both in paracrine and endocrine modes [43], accordingly, our transwell assay demonstrated that HGF can induce NEPHRIN and NEPH1 activation in an endocrine fashion (**Fig. 3B**). While the systemic deletion of HGF in mice induces early embryonic lethality [44, 45], the podocyte-specific knockout of its receptor c-MET did not result in any structural or physiological loss of renal function [46]. Importantly, we did not find any involvement of MET receptor in HGF-induced NEPHRIN and NEPH1 phosphorylation.

To conclusively establish that HGF can induce NEPHRIN and NEPH1 in a ligand-based fashion, we demonstrated a direct interaction of HGF with NEPHRIN and NEPH1. Furthermore, we showed that HGF interacted at specific sites in the extracellular IgG domain 2 of NEPH1 and IgG domain 3 of NEPHRIN, which are highly conserved. Interestingly, its binding affinity to NEPHRIN was more than 20-fold than that of NEPH1, which is comparable to the reported affinity of HGF for MET receptor (K_D_=0.2 – 0.3 nM [47]. How these different affinities affect the physiological role of this interaction will require further investigation. Since, HGF has renoprotective properties [48], we hypothesized that HGF induced renoprotection may involve its binding and activation of NEPH1 and NEPHRIN. This hypothesis was tested in two models of podocyte injury, where PS treatment of cultured human podocytes or Drosophila nephrocytes induced actin cytoskeleton disorganization [49] and loss of Sns (Sticks and stones) [50] respectively. Expectedly, exogenous addition of purified HGF to injured podocytes or nephrocytes induced recovery from injury, but more importantly, these recoveries were significantly attenuated by addition of inhibitory NEPHRIN or NEPH1 peptides that interacted with HGF. Collectively, these results highlight a mechanism, where renoprotective signals in podocytes are propagated via HGF induced activation of NEPHRIN and NEPH1.

Unlike the MET receptor tyrosine kinase [51], NEPHRIN and NEPH1 do not have a kinase activity and therefore, the question remains, how HGF binding results in the phosphorylation of these proteins. Although speculative, it is likely that HGF binding induces a conformational change in NEPHRIN and NEPH1 that results in the recruitment of SFK FYN at the intracellular region leading to their phosphorylation. To further understand these biophysical changes, structural analysis of these interactions using X-ray or SAXS-based measurement will be necessary and will be the subject of our future investigation.

## Methods

### Materials (Plasmids, antibodies and reagents)

Full-length Mus musculus NEPHRIN and NEPH1 cDNAs were cloned using standard PCR cloning procedure as described previously [22, 52]. HGF expressing plasmid pCMV3-Flag-HGF was procured from Sino Biological Inc. Substrate trapping SHP-2^DM^ (D425A/C459S) was provided Dr. Matt. [53] Purified human recombinant HGF, VEGF, TNFα were procured from Sigma. Polyclonal purified antibodies to NEPH1, phospho-NEPH1 and NEPHRIN have been previously described [24, 54]. GAPDH and anti-FLAG antibodies and Puromycin were procured commercially from Sigma-Aldrich. Antibodies against SHP-2, Phospho-NEPHRIN, MET receptor and FYN were obtained from Cell Signaling Technologies (Danvers, MA). NEPHRIN antibody was procured from Abcam. The SNS antibody is a gift from Dr. MP Krahn [55]. The cell transfection reagent Lipofectamine 2000 was purchased from Invitrogen (Cat. No. 11668019).

### Generation of lentiviral expression constructs for FLAG-NEPHRIN and FLAG-NEPH1

Standard PCR cloning technique was used to generate mammalian expression plasmid encoding FLAG-tagged full-length NEPHRIN and NEPH1. PCR amplified FLAG-NEPHRIN and NEPH1 were cloned at the EcoRI and EcoRI-Sal1 sites in the retroviral pBABE-puro vector respectively as described previously [25, 56]. The FLAG tag was inserted between the signal peptide and the start of the extracellular domains of NEPHRIN and NEPH1. The fidelity of constructs was determined by restriction digestion and DNA sequencing. Retroviruses overexpressing FLAG-NEPH1 and FLAG-NEPHRIN were generated by the transfection of the respective plasmids into Phoenix cells according to the manufacturer’s instructions. The cultured podocytes and HEK293 cells were transfected with these viruses and the expression of FLAG-NEPHRIN and FLAG-NEPH1 was evaluated by western blotting. Transfected cells were grown in 2.5μg/ml of puromycin-containing medium for the selection of stable transfectants. Details for the procedure of producing retrovirus and transfecting podocytes to generate a stable cell line have been described previously [22].

### Cloning, expression and purification of recombinant FLAG-NEPH1 full length (FL) and FLAG-NEPHRIN-extracellular domain (ECD)

To study the *in vitro* biochemistry and the kinetics of the interactions of NEPH1 and NEPHRIN with HGF, Baculoviral expression system was used. Bac to Bac Baculoviral Expression system (Invitrogen, Catalog nos. 10359-016) employing SF9 insect cell line was established for expressing the FLAG-NEPH1-FL and FLAG-NEPHRIN-ECD. The recombinant viral vector containing the gene insert of NEPH1 or NEPHRIN (rBacmid) was produced by transposition events in *E. coli* (DH10) host strain and blue white selection were generated and white colonies were picked. The insertion was confirmed by PCR and DNA sequencing. After confirmation, the rBacmid was transfected into SF9 cells and the viral particles were used to infect SF9 cells for protein production. Cells were harvested to purify HIS-FLAG-NEPH1-FL protein and HIS-FLAG-NEPHRIN ECD. The proteins were purified using Ni-NTA affinity column. Column bound proteins were eluted using 200 mM Imidazole on AKTA FPLC system as described previously [25].

### Cell culture growth and treatments

The immortalized human podocyte cell line was obtained from Dr. Moin Saleem [23]. and were cultured in RPMI 1640–based medium supplemented with 10% fetal bovine serum (Invitrogen), 2g/l of sodium bicarbonate (NaHCO3), insulin-transferrin-selenium supplement (Sigma-Aldrich, St. Louis, MO), and 200 units/ml penicillin and streptomycin (Roche Applied Science, Penzberg, Germany) as described previously)[54, 57]. HEK293 cells were cultured in Dulbecco’s modified Eagle’s medium supplemented with 10% fetal bovine serum (FBS; Invitrogen) and 200 U/ml of penicillin and streptomycin (Invitrogen). Lipofectamine 2000 (Invitrogen) was used to perform transfection according to the manufacturer’s protocol. HEK293 cells overexpressing FLAG NEPH1/NEPHRIN were plated and grown to 90% confluency and then serum starved for 2 hours. Cells were then treated with fresh pervanadate (1 mM) for 45 minutes at 37°C as described previously [58]. Vehicle treated cells were used as controls. Post pervanadate treatment, the cells were lysed in RIPA buffer supplemented with phosphatase and protease inhibitors. Protein estimation of each lysate was performed using the BCA protein estimation kit and equivalent amounts of lysates were subjected to SDS-PAGE and immunoblot analysis.

### *In vitro* phosphatase assays

To evaluate dephosphorylation of NEPHRIN and NEPH1 by SHP-2, GST-NEPH1-CD (cytoplasmic domain) or GST-NEPHRIN-CD [59] (2µg each) were mixed with purified active HIS-FYN (0.5 µg) immobilized on Ni-NTA beads in the presence of 1 mM ATP in 1X Kinase Buffer (25 mM HEPES at pH 7.4, 5 mM MgCl2, 5 mM MnCl2, 100 mM NaCl, 1 mM ATP) for 30 min at 30 C. The beads were removed by centrifugation and the phosphorylated GST-NEPHRIN and GST-NEPH1 were mixed with recombinant purified active SHP-2 (Recombinant Human SHP-2; Cat. No.: 1894-SH; R&D Systems) in the presence of phosphatase buffer (10mM HEPES, pH7.4, 0.1 mM EDTA and 1 mM DTT) and incubated at 37 C for 45-60 min. Reactions were stopped by adding sample loading buffer (Thermo Scientific, Cat. No. 39000) and heating at 95 °C for 5 min. The tyrosine phosphorylation of substrate proteins was analyzed by immunoblotting using anti-phospho NEPH1/NEPHRIN antibodies.

### Coimmunoprecipitation

Coimmunoprecipitations were performed as described previously with minor modifications (10). HEK 293T cells stably expressing FLAG-NEPH1 or FLAG-NEPHRIN were transiently transfected with HGF expressing pCMV-HGF plasmid using the Lipofectamine 2000 transfection agent following manufacturer’s instructions. After 48 h of incubation, cells were washed twice with PBS and lysed in RIPA buffer (phosphate-buffered saline [PBS] containing 0.1% sodium dodecyl sulfate [SDS], 1% Nonidet P-40, 0.5% sodium deoxycholate, and 100 mM potassium iodide) with EDTA-free proteinase inhibitor mixture (Roche Molecular Biochemicals). Lysates were cleared by centrifugation at 10000 rpm for 10 min at 4°C. After centrifugation, cell lysates containing equal amounts of total protein were incubated with anti HGF antibody or mice/rabbit IgG for 2 h at 4°C, followed by addition of 5µl (packed volume) of protein G-coupled Agarose beads (ROCHE) and continued overnight incubation at 4°C. Beads were then collected by centrifugation at 3000 rpm for 5 min at 4°C, extensively washed with PBST and resuspended in SDS gel loading buffer. The proteins were separated on a 10% SDS-polyacrylamide gel, transferred to a PVDF membrane, and analyzed by immunoblotting with the corresponding antibodies. Control samples were incubated with a nonspecific rabbit antiserum followed by the addition of protein G beads.

### Molecular modeling for predicting HGF binding regions in NEPH1 and NEPHRIN

The MET region involved in high affinity binding to HGF-α chain is located in the immunoglobulin-like domains IPT 3 and 4 of MET receptor [60]. For predicting the HGF binding site in NEPH1 and NEPHRIN, sequences corresponding to IPT 3 and 4 of MET receptor were separately aligned to the sequences of NEPH1 and NEPHRIN using the sequence alignment tool BLAST (pBlast). Sequences with highest identify and lowest gap were predicted to have high HGF binding probability. Two HGF binding regions for NEPHRIN from the IgG domain 2 PDITILLSGQTISDISANVNEGSQQKL (NEPHRIN-P1) and: FTVEATARVTPRSSDNRQLLVCEASS (NEPHRIN-P2); and one region from the NEPH1 IgG 3 domain: QEGERVVFTCQATANPEIL (NEPH-P1) were predicted. To further confirm the binding region, online server for protein-protein docking Z dock was used. Extracellular domains of NEPH1 and NEPHRIN, and HGF-α chain were modeled using protein fold recognition-based modeling server PHYRE2 as previously described [61]. PHYRE2 used following PDB templates to generate the models: for HGF, the template used was 4dur.2.A, x-ray crystal structure of full-length type ii human plasminogen (93% coverage, 100% confidence), for NEPHRIN it was 3b43A, X-ray crystal structure of titin (89% coverage, 100% confidence), for NEPH1 it was 3b43A, X-ray crystal structure titin (85% coverage, 100% confidence), and for MET it was 5L5C, the X-Ray crystal structure of Plexin A1 full extracellular region (81% coverage, 100% confidence). The modeled proteins were used in the online protein-protein docking server (Z dock). Predicted model for MET-HGF (α-chain), NEPHRIN-HGF(α-chain) and the NEPH1-HGF (α-chain) was found to bind with equivalent energies in the predicted regions.

### Recombinant proteins and Peptides

Proteins, including glutathione S-transferase (GST)-NEPH1 cytoplasmic domain (CD) and GST-NEPHRIN CD were expressed and purified from Escherichia coli BL21 cells (Stratagene). Purified phosphorylated GST-NEPH1 was expressed and purified from TKB1 cells (Stratagene). Details of the expression and purification protocol have been described previously [59, 62]. The predicted NEPHRIN and NEPH1 peptide sequences (NEPHRIN-P1, P2 and NEPH1-P1) described above were synthesized chemically using standard 9-fluorenylmethoxy carbonyl (Fmoc) solid-phase chemistry, and 2-chloro trityl resin was used as a solid support (PTI peptide synthesizer). Post synthesis, the peptides were cleaved using a cocktail of trifluoroacetic acid (TFA) and scavengers and were purified to homogeneity using high-performance liquid chromatography (HPLC) (C18 semi prep and analytical columns attached to a Waters system). Identities of the purified peptides were confirmed by MALDI-TOF MS (Voyager).

### Dot Blot assay

The NEPH1-P1 and NEPHRIN-P1 and P2 peptides were used in a dot blot assay as described previously [63]. Briefly, the test and control peptides (300 ng) were spot blotted in duplicate onto nitrocellulose membrane and allowed to air dry. Scrambled peptide was used as a negative control and purified recombinant HGF was used as a positive control. The membrane was blocked for 1h using skim milk and then incubated with HGF (1µg recombinant purified HGF in 5% BSA in PBS) at 4°C and then immunoblotted with anti HGF antibody.

### Surface plasmon resonance (SPR)

The real-time binding experiment was performed at BMISR (Biacore Molecular Interaction Shared Resource) center using the Biacore T200 instrument. HGF was immobilized on the CM5 chip cell, and the instrument was programmed to perform a series of binding assays with increasing concentrations of FLAG-NEPH1-FL and FLAG-NEPHRIN-ECD. Surface regeneration and complete dissociation between the two proteins were achieved by using 2M NaCl. 1:1 kinetics model fitting was used for the analysis of sensorgrams and calculating the kinetic constants (association constant [Ka], dissociation constant [Kd], and equilibrium dissociation constant [K_D_].

### HGF influx assay using Transwell co-culture plates

HEK293 cells overexpressing HGF were cultured in the upper chamber of a 6 well cell culture transwell filters plate (0.4-μm pore; Corning) and in the lower chamber, FLAG-NEPH1 or FLAG-NEPHRIN overexpressing HEK293 cells were cultured. The control upper chamber consisted of non-HGF expressing HEK293 cells. The cells in the lower chamber were harvested after 48h, lysed and their phosphorylation was assessed by western blotting using phospho-antibodies for NEPH1 and NEPHRIN.

### Immunofluorescence microscopy

Podocytes stably expressing FLAG-NEPHRIN and FLAG-NEPH1 were plated and grown to 80-90% confluency on glass coverslips coated with collagen. The cells were then serum starved over-night, washed with PBS and subjected to injury by incubating with 500µg/ml of Protamine Sulfate for 8 h at 33 °C in serum free media. Cells were then incubated with and without recombinant human active HGF (cell culture grade, 50ng/ml) in 2% serum containing RPMI media for 4, 8 and 12 hrs. at 33 °C. Control sets included, no treatment, NEPH1 Knockdown podocytes or podocytes where co-incubation with NEPH1 or NEPHRIN inhibitory peptides (HGF binding peptides) was performed. The cells were washed with PBS and fixed with 4% paraformaldehyde (in 1 × PBS), followed by permeabilization with 0.1% SDS. Immunostaining was performed using fluorescent labeled Phalloidin (Alexa Flour 488) in dark at room temperature for 1 hr. After 4 washes with PBS, the coverslips were mounted with GelMount containing DAPI to label nuclei and images were collected the following day. Fluorescence microscopy was performed using Leica Confocal microscope (TCS, SP5 model). All parameters were kept constant (including exposure time) while taking images. Images were taken in 16 bits format and were analyzed using ImageJ software from single-plane images. Brightness and contrast adjustments were kept constant throughout the images. Representative images from three independent experiments are shown.

### ShRNA-mediated Knockdown of MET receptor, NEPH1 and SHP-2 in cultured podocytes

MET and NEPH1 knockdown in cultured human podocytes was achieved using MET and NEPH1 specific shRNA. The shRNAs in pLKO.1-puro (MET MISSION Lentiviral Transduction shRNA particle) were commercially purchased from Sigma (Cat. No. SHCLNV-NM_000245 and TRC number: TRCN0000196443 for Met; and SHCLNV-NM_018240 and TRC number: TRCN0000147545 for NEPH1). SHP-2 knockdown in podocytes was generated as described in previously [53]. Transduction of the shRNA plasmids was performed according to the manufacturer’s instructions. Selection for stably transduced cells was performed by culturing the cells in 2.5μg/ml puromycin-containing medium, and the MET receptor and NEPH1 knockdown was confirmed by Western blot.

### Drosophila nephrocytes as a model for HGF induced renoprotection

The Drosophila experiments were performed at the University of Maryland under the guidance of Dr. Zie Han. Briefly, Drosophila pericardial nephrocytes were dissected in artificial hemolymph and then treated under different conditions for 90 minutes at room temperature. The final concentration of PS was 1mg/ml, HGF 50ng/ml, NEPH1 peptide 250µg/ml, NEPHRIN P1 25µg/ml, NEPHRIN P2 25ug/ml. The nephrocytes then were heat fixed for 20 seconds and stained with anti-SNS antibody. For quantification, 7 nephrocytes were analyzed from three flies for each treatment.

### Statistical analysis

All statistical analyses were performed using GraphPad Prism 7 software. Each data set is presented as mean ± SEM. The Mann-Whitney (nonparametric) test was performed to assess the statistical differences between 2 groups; to analyze differences between more than 2 groups, a 1-way (Kruskal-Wallis test) or 2-way (Sidak’s multiple comparison test) or Tukey’s multiple comparisons test (One-way anova) nonparametric analysis of variance was performed. Details of the statistical analyses used for each experiment are provided in the respective figure legends. A P of ≤0.05 was considered statistically significant.

## DATA AVAILABILITY

The authors declare that all data supporting the finding of this study are available within this article and its supplementary information files or from the corresponding author upon request. The plasmids will be available from the corresponding author upon reasonable request.

## DISCLOSURE

All the authors declared no competing interests.

## ACKNOWLEDGMENTS

This work was supported in whole or in part by NIH grants 2R01DK087956-06A1 and R56 DK116887-01A1 to DN.

## AUTHOR CONTRIBUTIONS

AKS, PS, EA, BR, CMF, AS and PW conducted the experiments and analyzed data. SKH, ZH, ML, JHL, GL, WRF helped with experimental designs and provided critical reagents. DN, EA and AKS designed the experiments, interpreted data, and wrote the manuscript. All the authors discussed results and commented on the manuscript

## Supplementary Figure legends

**Supplementary Figure 1:**
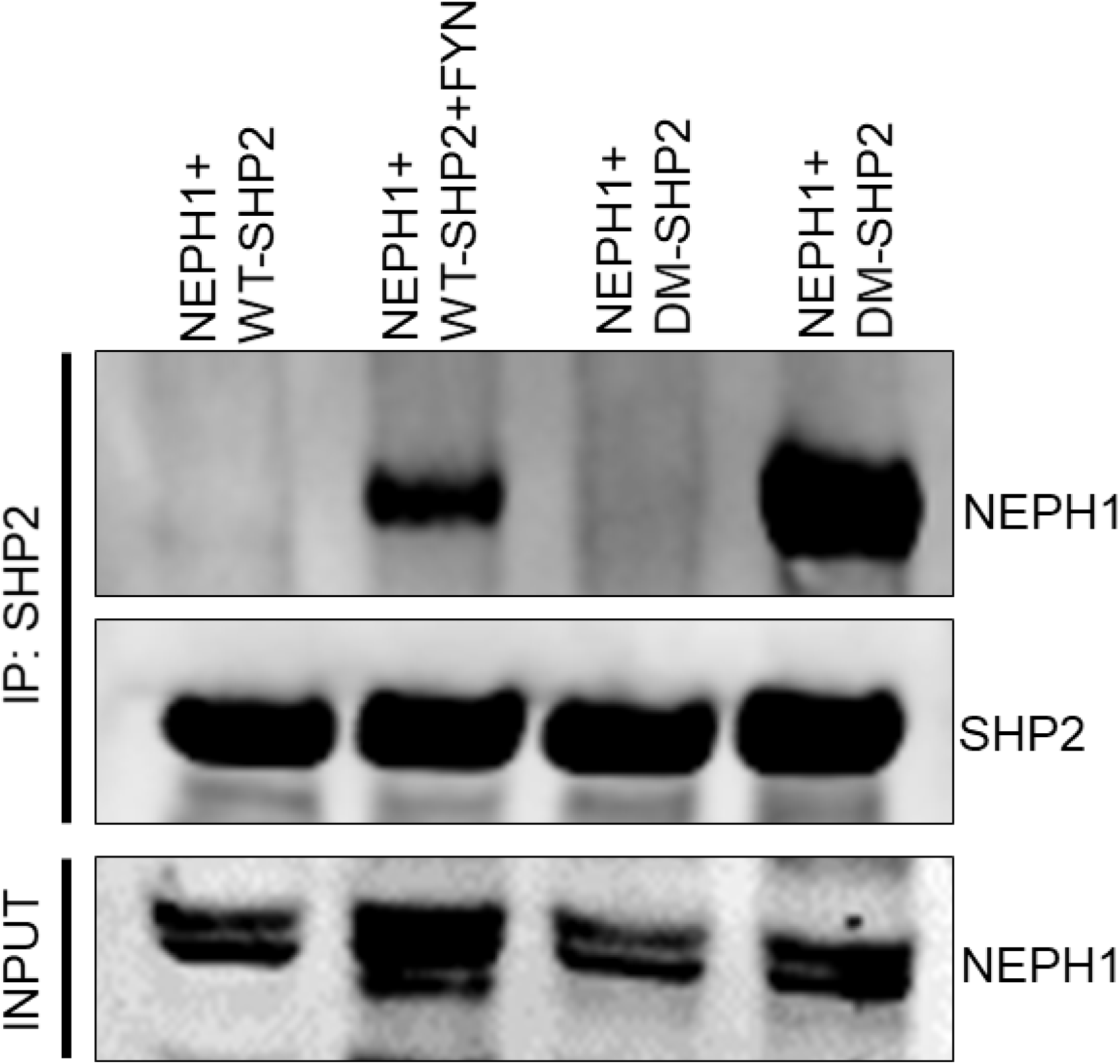
NEPH1 was coexpressed with SHP-2 or the substrate trapping SHP-2 mutant (DM SHP-2) without or with FYN. To evaluate NEPH1 binding, SHP-2 was immunoprecipitated form the cell lysate and western blotted with NEPH1 antibody, which NEPH1 bound SHP-2 in the presence of FYN and significantly enhanced binding was observed with DM SHP-2 indicating NEPH1 as a SHP-2 substrate.

**Supplementary Figure 2:**
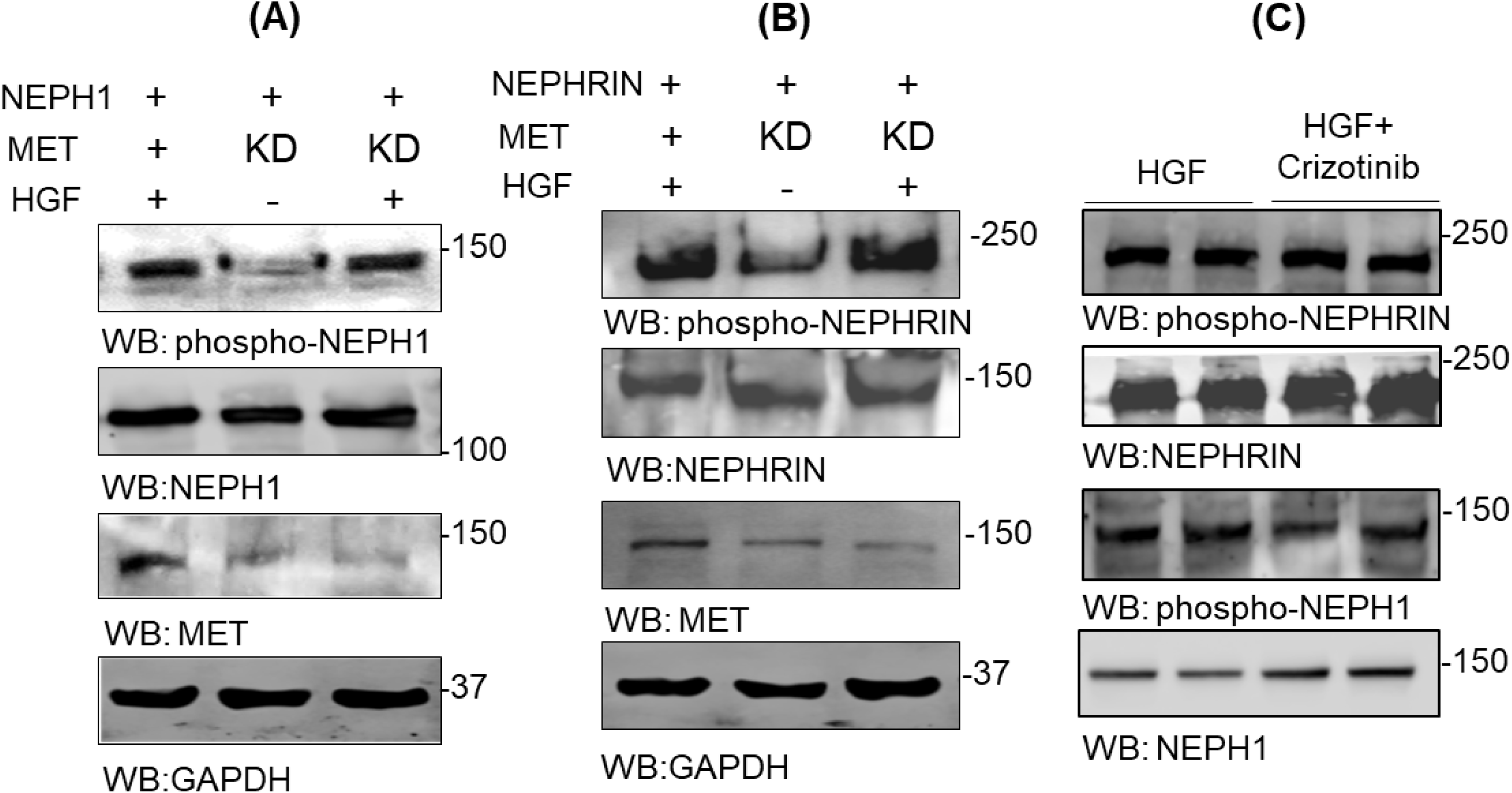
MET receptor is not required for HGF-induced does not NEPHRIN and NEPH1 phosphorylation: **(A & B)** FLAG-NEPH1 or FLAG-NEPHRIN were overexpressed in HEK293 cells with stable shRNA-mediated knockdown (KD) of MET receptor. Recombinant active HGF **(20ng/ml)** was exogenously added to these cells and NEPHRIN and NEPH1 phosphorylation in cell lysate was evaluated, which showed no change in their phosphorylation due to the loss of MET receptor. **(C)** Recombinant active HGF **(20ng/ml)** was exogenously added to the HEK293 cells overexpressing FLAG-NEPH1 or FLAG-NEPHRIN in the absence or presence of 100nM Crizotinib (MET receptor inhibitor) and the phosphorylation of NEPH1 and NEPHRIN were analyzed. The inhibitor had no effect on NEPH1 and NEPHRIN phosphorylation.

**Supplementary Figure 3:**
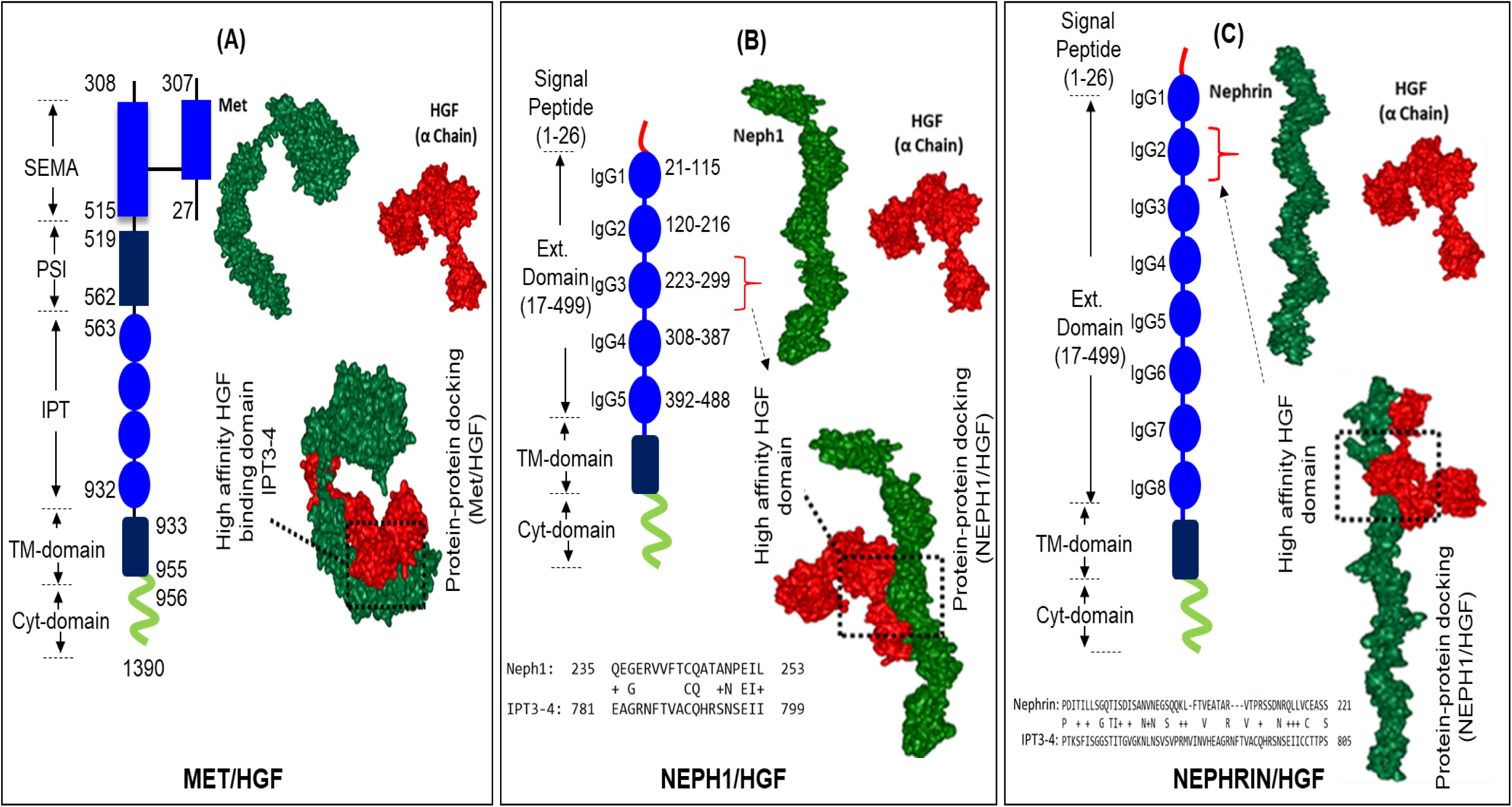
Identification of HGF interacting domains in NEPHRIN and NEPH1 through molecular modeling analysis. **(A-C)** The high affinity binding site of MET receptor (IPT3-4 domain) with HGF was used as a template. The cartoon represents the respective domain architecture of MET (**A**), NEPH1 (**B**), and NEPHRIN (**C**). The alignment of protein sequences representing the extracellular domains of NEPH1 and NEPHRIN with IPT3-4 region of MET was performed using NCBI BLAST, which showed 32% and 23% identities with IgG domain 3 of NEPH1 and IgG domain 2 of NEPHRIN respectively. Peptides from these putative HGF binding sites were synthesized and used in binding and functional assays. Structural protein models for the HGF α chain, MET, NEPH1 and NEPHRIN extracellular domains were constructed using protein fold recognition-based modeling server PHYRE2. Using Z dock protein-protein docking with HGF α chain was performed that identified the interacting regions.

**Supplementary Figure 4:**
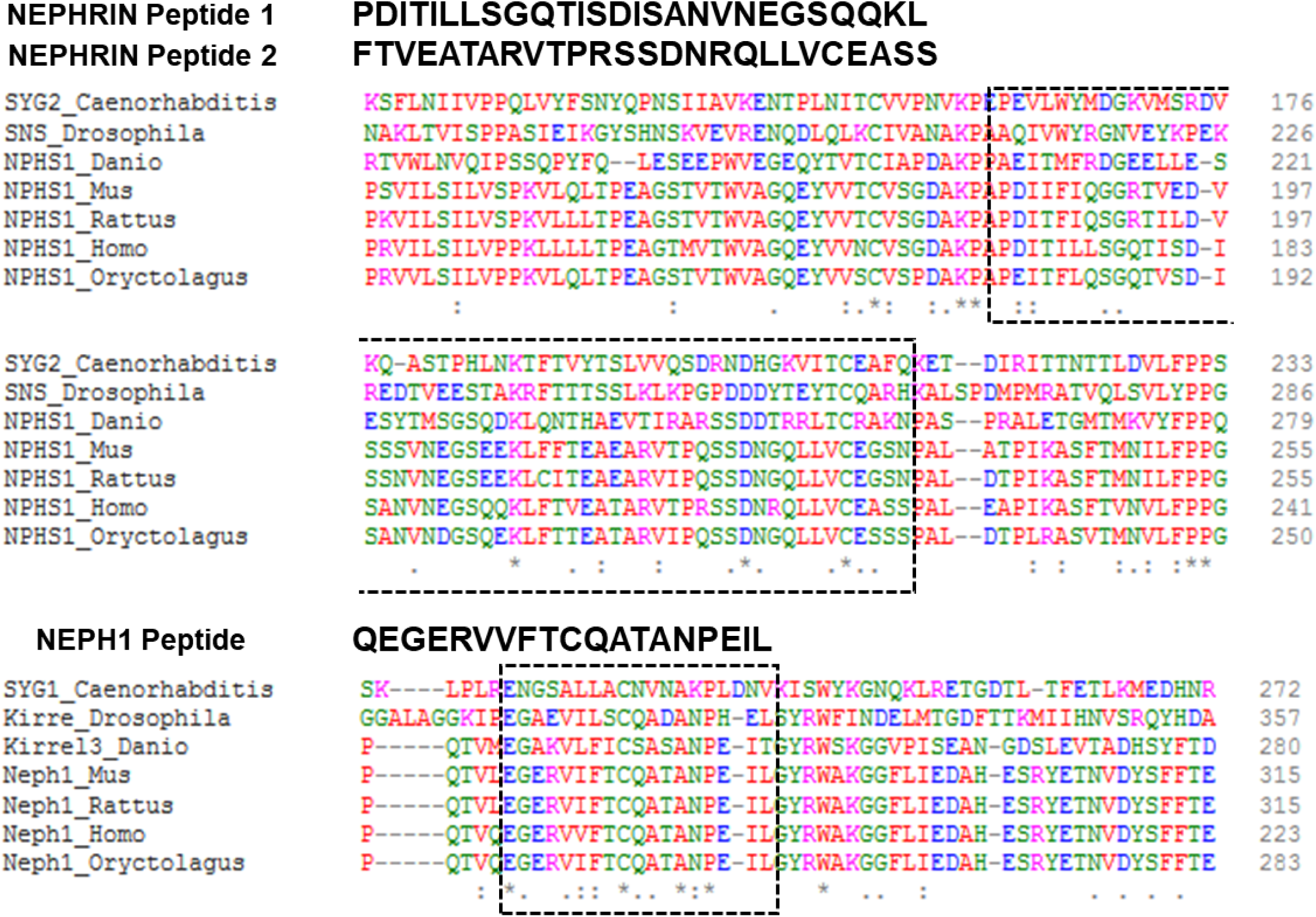
HGF binding region in NEPHRIN and NEPH1 is highly conserved: The NEPHRIN (A) and NEPH1 (B) peptides that interacted with HGF were highly conserved across several species. including rat, mice, human, C elegans, zebrafish and rabbit.

**Supplementary Figure 5:**
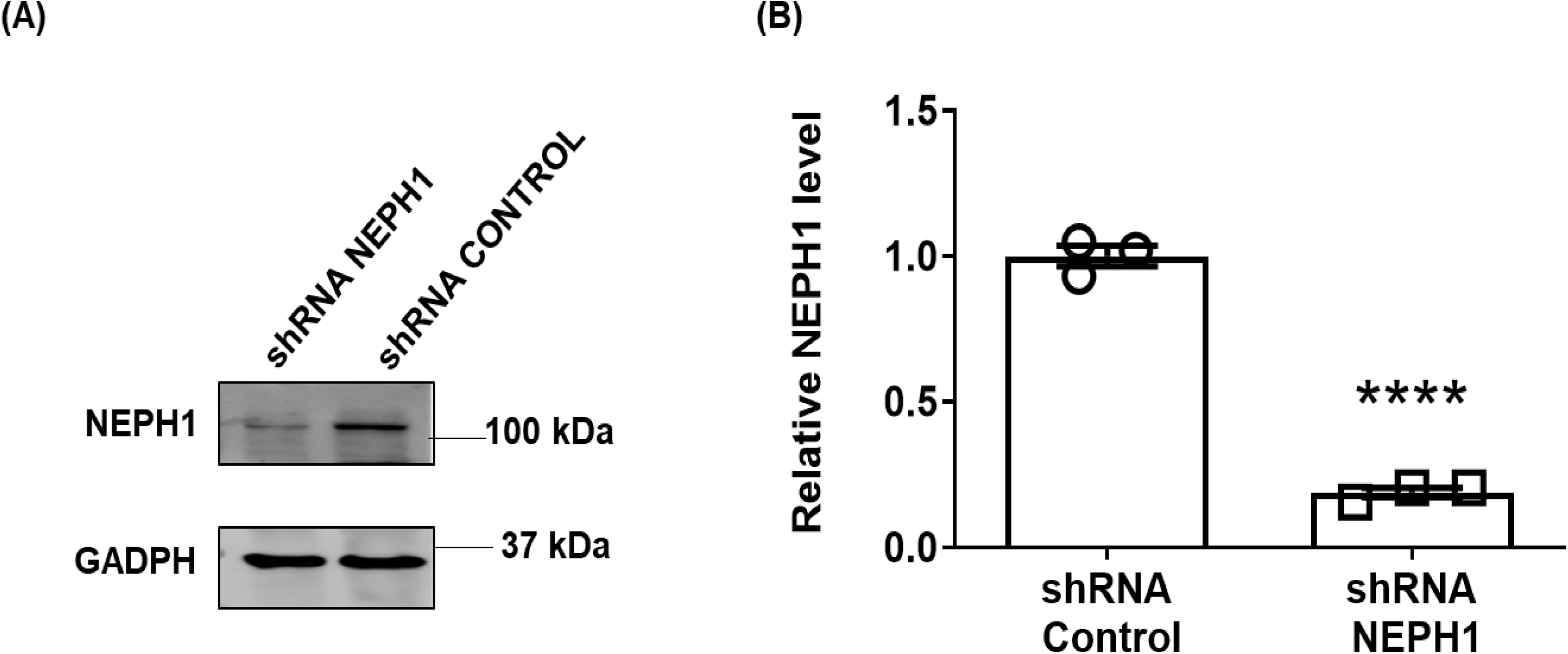
NEPH1 knockdown in cultured human podocytes. (**A**) Podocytes with stable NEPH1 knock down were prepared using NEPH1-specific sh-RNA. NEPH1 knockdown was evaluated by western blotting with NEPH1 antibody GAPDH was used to determine equivalent protein loading control (**B**) The relative decrease in NEPH1 levels were quantified. All experiments were performed in triplicate. Data are presented as mean ± SEM, and P values were calculated using two-tailed student’s t test. ****P ≤ 0.0001

**Supplementary Figure 6:**
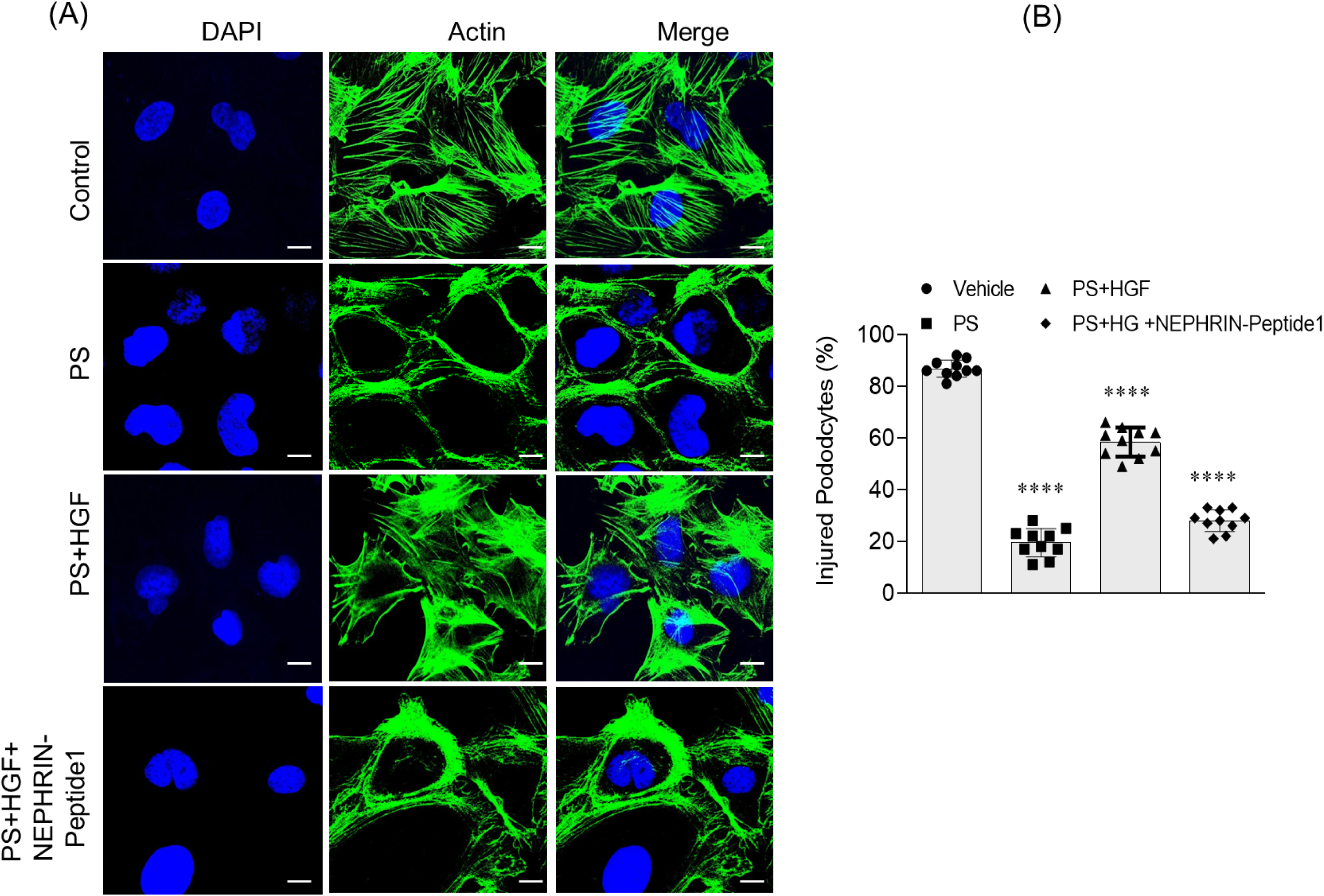
HGF treatment repairs podocytes/nephrocytes in a NEPH1/NEPHRIN dependent fashion: (**A & B**) Cultured human podocytes overexpressing NEPHRIN were treated with PS and actin cytoskeleton (green) disorganization was visualized by phalloidin staining. To induce recovery, HGF (50 ng/ml) was added to the PS treated podocytes. Addition of NEPHRIN inhibitory peptide blocked HGF-induced recovery. 10 cells per experimental condition were evaluated from three experimental repeats. Scalebar=25µm. Data are presented as mean ± SEM, and P values were calculated using the Tukey’s multiple comparisons test (One-way Anova). ****P ≤0.0001

**Supplementary Figure 7:**
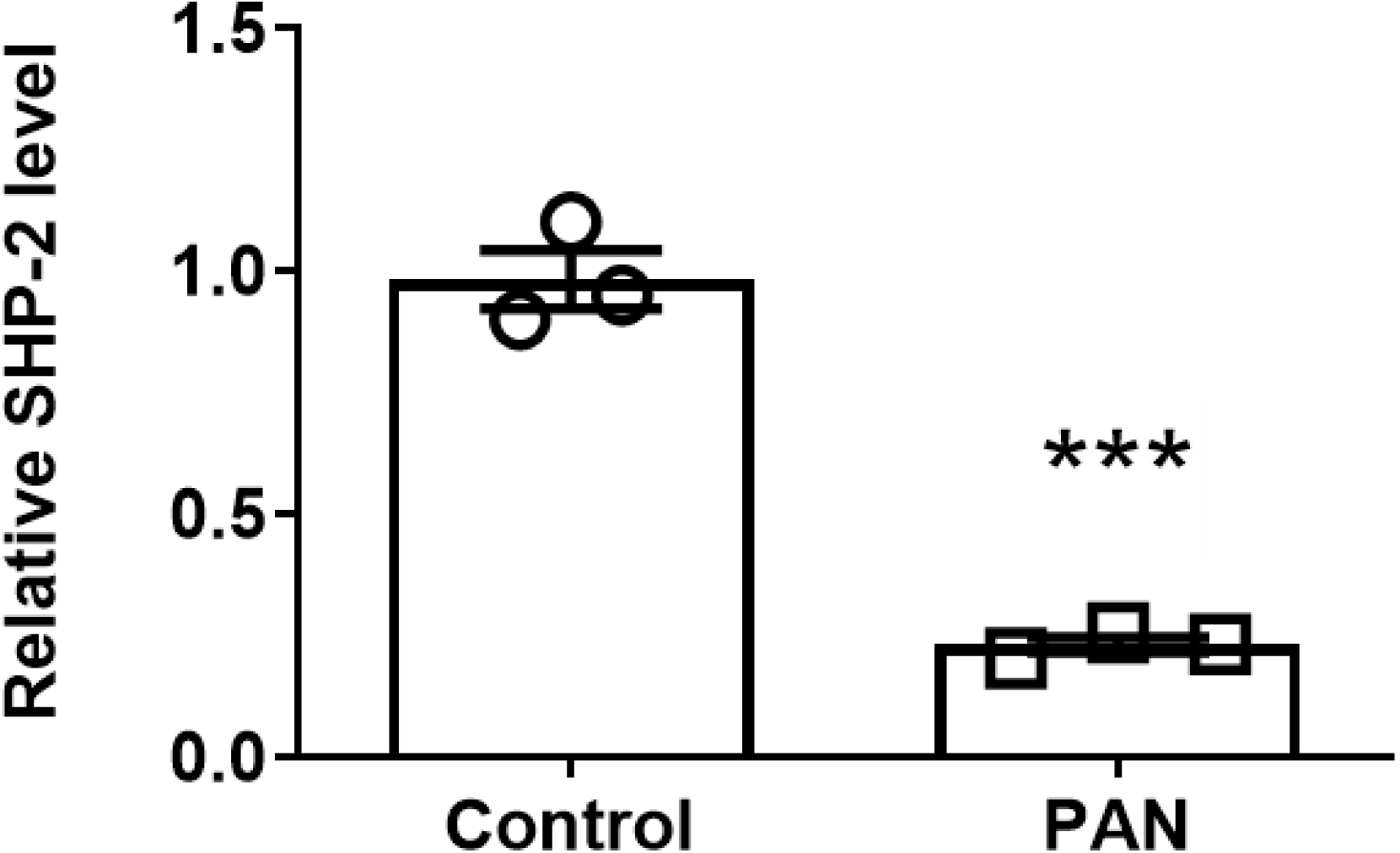
Quantification of the SHP-2 expression level in PAN injured podocytes: The relative SHP-2 expression measured from the mRNA profiling data of podocytes injured with PAN is presented and shows down-regulation of SHP-2 expression. All experiments were performed in triplicate. Data are presented as mean ± SEM, and P values were calculated using two-tailed student’s t test. ***P ≤ 0.001

**TABLE 1a:**
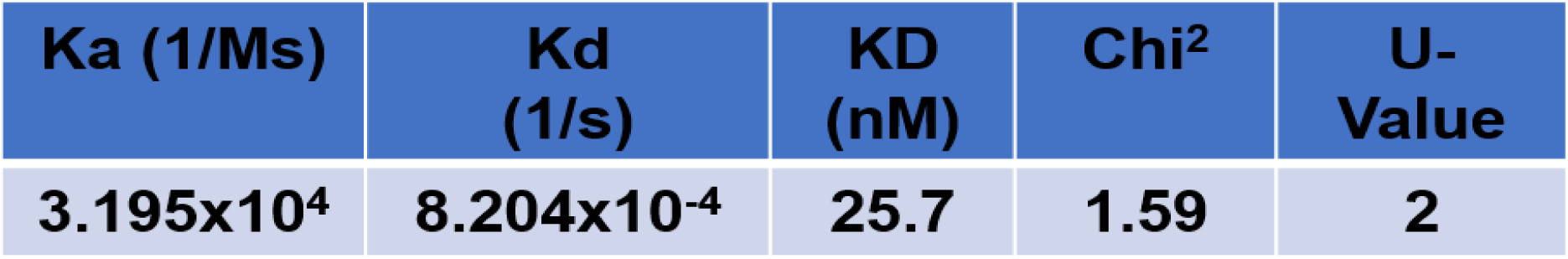
Summary of the results obtained from SPR analysis for NEPH1.

**TABLE 1b:** Summary of the results obtained from SPR analysis for NEPHRIN-ECD.

| Ka (1/Ms) | Kd (1/s) | KD (nM) | Chi<sup>2</sup> | U-Value |
| --- | --- | --- | --- | --- |
| 2.066x10<sup>6</sup> | 7.258x10<sup>-4</sup> | 0.4 | 6.18 | 4 |

## Notes

### Competing Interest Statement

The authors have declared no competing interest.

